# Style Transfer Generative Adversarial Networks to Harmonize Multi-Site MRI to a Single Reference Image to Avoid Over-Correction

**DOI:** 10.1101/2022.09.12.506445

**Authors:** Mengting Liu, Alyssa H. Zhu, Piyush Maiti, Sophia I. Thomopoulos, Shruti Gadewar, Yaqiong Chai, Hosung Kim, Neda Jahanshad, the Alzheimer’s Disease Neuroimaging Initiative

## Abstract

Recent work within neuroimaging consortia have aimed to identify reproducible, and often subtle, brain signatures of psychiatric or neurological conditions. To allow for high-powered brain imaging analyses, it is often necessary to pool MR images that were acquired with different protocols across multiple scanners. Current retrospective harmonization techniques have shown promise in removing cross-site image variation. However, most statistical approaches may over-correct for technical, scanning-related, variation as they cannot distinguish between confounded image-acquisition based variability and cross-site population variability. Such statistical methods often require that datasets contain subjects or patient groups with similar clinical or demographic information to isolate the acquisition-based variability. To overcome this limitation, we consider cross-site MRI image harmonization as a style transfer problem rather than a domain transfer problem. Using a fully unsupervised deep-learning framework based on a generative adversarial network (GAN), we show that MR images can be harmonized by inserting the style information encoded from a single reference image, without knowing their site/scanner labels *a priori*. We trained our model using data from five large-scale multi-site datasets with varied demographics. Results demonstrated that our style-encoding model can harmonize MR images, and match intensity profiles, without relying on traveling subjects. This model also avoids the need to control for clinical, diagnostic, or demographic information. We highlight the effectiveness of our method for clinical research by comparing extracted cortical and subcortical features, brain-age estimates, and case-control effect sizes before and after the harmonization. We showed that our harmonization removed the cross-site variances, while preserving the anatomical information and clinical meaningful patterns. We further demonstrated that with a diverse training set, our method successfully harmonized MR images collected from unseen scanners and protocols, suggesting a promising novel tool for ongoing collaborative studies. Source code is released in USC-IGC/style_transfer_harmonization (github.com).

## 1. Introduction

Neuroimaging studies often need to collect data across multiple sites to ensure the sample size and power required to obtain reliable and robust results. Combining multi-site MRI data, however, is a non-trivial issue as images are subject to both acquisition- and cohort-based variability. Acquisition-based variability is often due to scanning factors such as manufacturer, magnetic field strength, coil type and number of channels and parameters such as image resolution and those related to the pulse sequence. Even when imaging parameters are prospectively planned to be as consistent as possible across scanning sites, the need for retrospective harmonization is often inevitable. Scanners in long-running studies such as the Alzheimer’s Disease Neuroimaging Initiative (ADNI), a multisite initiative that has been ongoing for nearly two decades [1], may undergo scanner drift [2], or scanner or software upgrades that alter image contrast or signal to noise ratios. International neuroimaging consortia have shown the need for effective retrospective neuroimaging harmonization methods. For example, the ENIGMA consortium has pooled information from hundreds of data collection sites around the world for collaborative initiatives aimed at identifying brain signatures of psychiatric or neurological conditions, charting brain trajectories, and even mapping the genetic architecture of brain structure. While initial efforts focused on harmonizing analytical plans towards coordinated meta-analyses, pooling individal-level data allows researchers to pose more targeted questions about factors that may not be similarly distributed across participating sites such as disease staging, or infrequent conditions, such as clinical diagnoses for transdiagnostic studies of history of suicide attempts [3]. These population-based differences are another source of inter-site differences. Even with the data collected from the same scanners/sites, the images collected may also show slight variance [4].

Existing retrospective data harmonization techniques have shown promise in removing cross-site variance from different studies to allow for such pooled analyses. Most harmonization methods fall into two broad categories: 1) harmonization of image-derived features using statistical properties of the distribution, for example ComBat [5–7]; 2) harmonization of the latent features extracted from the MR images for a single cross-site specific task, such as a regional segmentation, disease classification, or age prediction. The main drawback of the first category of harmonization techniques is that it requires many statistical assumptions that may be difficult to satisfy. An extensive review of the first category of harmonization techniques was performed by Bayer et al., [8].The second category, which seeks to circumvent the statistical assumption pitfalls by avoiding the harmonization of datasets directly but focusing on the task output, is largely composed of deep learning-based approaches, namely domain adaptation techniques [9, 10]. Domain transfer learning and domain adversarial learning have been applied for MRI harmonization [11].

While task-specific harmonization can be powerful for a particular outcome, if a wide range of tasks were to be performed on images, harmonization would need to be performed separately for each task, resulting in inconsistent, task-dependent harmonization schemes [12]. There are also some applications, like cortical surface construction (as opposed to segmentation), that cannot be directly embedded into a deep learning framework [13]. Such cases require image translation or a direct image-to-image harmonization for MRIs. Several image translation-based harmonization methods have been proposed. Supervised methods typically require traveling subjects and must be planned prospectively [14]. Unsupervised methods, such as variational auto-encoders [15] or CycleGAN [5], often separate MR images into well-defined domains in terms of scanners or sites. These methods may be prone to overcorrection, by which we mean correcting for biological factors in addition to, or perhaps instead of, technical scanning-related variables. As each site gets a different label, two different sites with scanning protocols that are similar, and populations that are different, may inadvertently adjust for biological differences rather than scanner differences.

In addition to the common challenges in unsupervised harmonization, most existing domain-based harmonization algorithms may not generalize to previously unseen sites [16]. When there is a new dataset to be harmonized that was not included in the training set, the harmonization usually cannot perform well and re-training to include the new dataset is typically required. Domain-based harmonization approaches, therefore, lack flexibility. They restrict the image harmonization to a very limited number of groups with clear borders and any images beyond the scope of these borders may not adequately harmonize. This could limit the applicability of those harmonization methods in large-scale consortia settings where there are a large number of studies with relatively small sample sizes with diverse acquisition protocols that are iteratively being added to studies.

Recently, deep learning methods have successfully completed diverse image translations by disentangling the image into ‘content’ and ‘style’ spaces [17–19], where contents represent low level information in images like contours and orientations, and styles can be considered high level information such as colors and textures. Images within the same group share the same content space but may show different styles. In MR images, we can consider the biologically defined anatomical information as the content, and the non-biological information such as intensity variation, signal-to-noise ratio (SNR), and contrasts as styles. Dewey and colleagues [20] used this breakdown to show promising results for MRI harmonization when paired image modalities from the same subjects were available to supervise the extraction of the content information; unfortunately, the same two sets of paired images are not always available across multiple datasets. In other work, Jiang et al., [21] also used a similar framework to facilitate the cross-domain image translations, where each image modality was treated as a domain. The problem faced by [21] was that styles that span multiple domains must be modeled together using a variational auto-encoder with a universal prior.

Here, we also consider that every single image can be disentangled into content and style, and image harmonization is a pure style transfer problem rather than a domain transfer problem, which is presumed by conventional domain-based approaches. We consider anatomical patterns (or contents) from MR images collected from different sites to share the same latent space, such that it is not necessary to separate them into different “domains”. These style-irrelevant patterns can be learned using an unsupervised cycle consistency generative adversarial network (GAN) model, and thus, paired modalities or any other paired information are not needed from the same subjects. Due to scanner shifts, and software upgrades, the styles for all the images, even those collected from the same scanner may be different. We consider every single image as a unique “domain” with its own style, and the styles can be learned flexibly using an adversarial approach, instead of using a universal prior distribution as in [21]. Furthermore, inspired by [22], we proposed that the style information needed for harmonization can be encoded from a single reference MR image directly. In short, the whole harmonization procedure relies solely on a source image and a reference image, where the source image provides the content or the biological architecture of the brain, and the reference image provides the target style information or the image intensity distribution and contrast.

To illustrate the clinical effectiveness of our approach, we train our model on healthy subsets of five publicly available neuroimaging datasets, including: the UK Biobank (UKBB), Parkinson’s Progression Markers Initiative (PPMI), Alzheimer Disease Neuroimaging Initiative (ADNI), Adolescent Brain Cognitive Development (ABCD) and International Consortium for Brain Mapping (ICBM). We use automated software, specifically FreeSurfer, to extract several commonly used metrics of interest and compare the features extracted before and after the harmonization. We show that our harmonization method successfully removed the cross-site variances, while preserving the anatomical information as demonstrated by retaining case/control effect sizes before and after harmonization within a single site. Using unseen traveling subjects, we further illustrate that our model successfully harmonized MR images collected from unseen scanners and protocols, suggesting a promising novel tool for ongoing collaborative studies.

## 2. Materials and Methods

### 2.1. The Architecture of Style-Encoding GAN

Let *X* be a set of full brain MR image slices. Given an image *x* ∈ *X*, our goal is to train a single generator *G* that can generate diverse images that correspond to the image *x* with a style code *s*, where *s* is associated with the style (non-biological) patterns from another image. The style code *s* is generated by a mapping network *M* from sampling a given latent vector *z* (*s* = *M*(*z*)). [23] explain the rationale for using *s* instead of *z*. Then, the generator *G* translates an input image *x* into an output image *G*(*x, s*) that reflects the style of *s*. To validate that the style code *s* is successfully injected into the output image *G*(*x, s*), another style encoding network *E* was designed to encode the style of *s* from images. That is, given an image *x*, the encoder *E* extracts the style code *s* = *E*(*x*) of *x*. *E* can produce diverse style codes using different images. This allows *G* to synthesize an output image reflecting the style, *s*, from different reference images of *X*. The goal of the network is to train *E* so that *E*(*G*(*x, s*)) = *s*, meaning that if an image was generated based on style code *s*, then *s* can also be encoded when this image was input into the style encoder *E*. Adaptive instance normalization (AdIN) [24] was used to inject *s* into *G*. Finally, the discriminator *D* learns a binary classification determining whether an image is a real image or a fake image as produced by *G, G*(*x, s*). As with [25], our model includes only one generator, one discriminator, and one style encoder (**Figure 1**).

**Figure 1.**
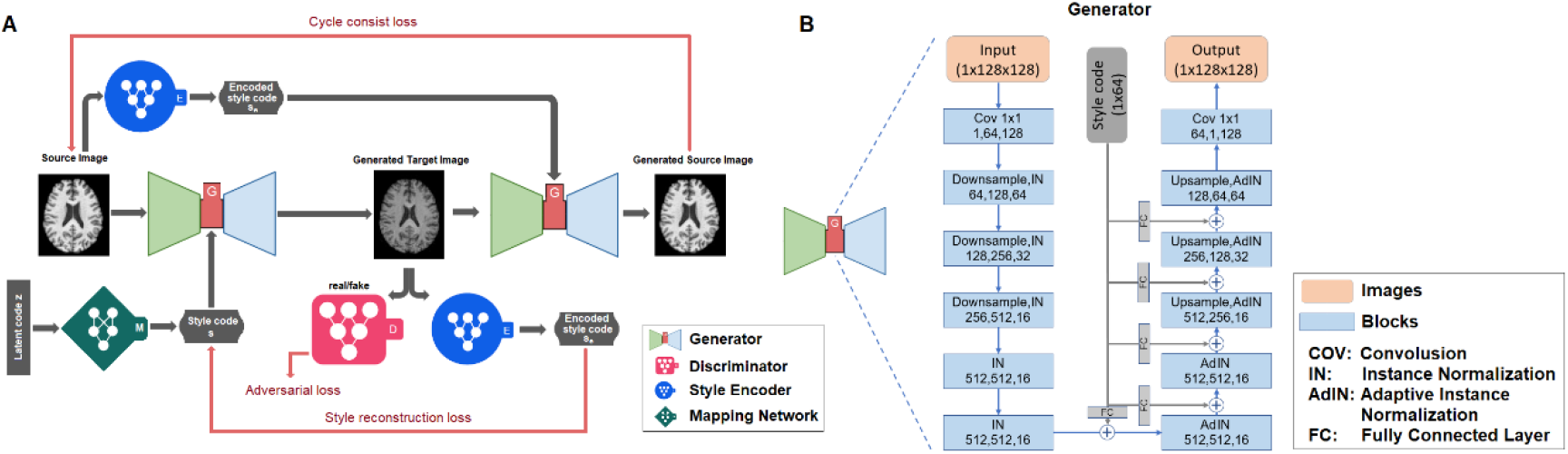
A) The architecture of the style-encoding GAN. The generator learns to generate an image by inputting a source image and a style code. B) The detailed architecture of the generator in the network. In each of the blocks in the process, the three numbers represent the number of input channels, number of output channels, and the image size.

#### 2.1.1 Network Training

Given an image *x* ∈ *X,* we train our framework using the following objectives: an adversarial loss; a cycle consistency loss; a style reconstruction loss; and a diversification loss, all of which we describe below.

- **Adversarial loss.** During training, we sample a latent code *z* ∈ *Z* randomly, and the mapping network *M* learns to generate a target style code *s* = *M* (*z*). The generator *G* takes an image *x* and style *s* as inputs and learns to generate an output image *G*(*x, s*) that is indistinguishable by the discriminator *D* from real images via an adversarial loss:

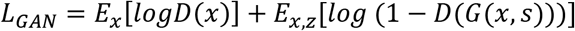
- **Cycle-consistency loss.** To guarantee that the generated images are consistent with the original images and properly preserving the style-irrelevant characteristics (e.g. anatomical patterns) of input *x*, an additional cycle consistency loss [26] is defined as the difference between original and reconstructed images:

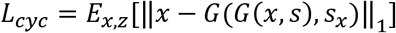

where *s_x_* = *E*(*x*) is the estimated style code of the input image *x*. By encouraging the generator *G* to reconstruct the input image *x* with the estimated style code *s_x_, G* learns to preserve the original characteristics of *x* while changing its style faithfully.
- **Style reconstruction loss.** In order to enforce the generator *G* to use the style code while generating the image *G*(*x, s*), we incorporate a style reconstruction loss:

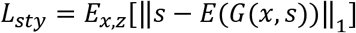 Our learned encoder *E* allows *G* to transform an input image *x*, to reflect the style of a reference image.
- **Style diversification loss.** To further enable the generator *G* to produce diverse images, we explicitly regularize *G* with the diversity sensitive loss [27]:

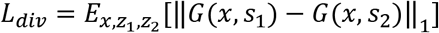

where the target style codes *s*_1_ and *s*_2_ are produced by *M* conditioned on two random latent codes *z*_1_ and *z*_2_ (i.e. *s_i_* = *M*(*z_i_*) for *i* ∈ [1]). Maximizing the regularization term forces *G* to explore the image space and discover meaningful style features to generate diverse images.

Put together, our full objective function can be summarized as follows:

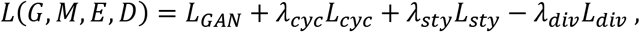

where *λ_cyc_, λ_sty_* and *λ_div_* are hyperparameters for each term. If the groups of images to be harmonized are confounded by demographic or clinical/pathological differences, such biological differences could also be inadvertently learned during the harmonization. To avoid this, we tuned the *λ_cyc_* in our model to preserve the style-irrelevant characteristics, including anatomical structure.

We set *λ_cyc_*=10 to make sure the biological content can be well preserved. We also set *λ_sty_*=1 and *λ_div_*=1. To stabilize the training, the weight *λ_div_* is linearly decayed to zero over the 200K iterations. We adopt the non-saturating adversarial loss [28] with R1 regularization [29] using γ = 1. We use the Adam [30] optimizer with *β*_1_ = 0 and *β*_2_ = 0.99. The learning rates for G, D, and E are set to 10^**−4**^, while that of M is set to 10^**−6**^.

#### 2.1.2 3D Image Reconstruction

Due to the GPU memory limitations, our network architecture has been designed for 2D images. This is in contrast to modern MRI scans which are 3D volumes [14]. We hence reconstruct the MRI 3D volumes by stacking the 2D slices. The model can be extended to a fully 3D deep network if GPU memory allows. Stacking 2D slices does not work well for slices at the more peripheral edges of the structure of interest, in this case, the brain; these slices contain fewer brain tissue types/contrasts to help ensure the model learns style features properly. To balance the GPU memory limitation and the edge-slice effect, we applied the harmonization on partial-3D-volumes, stacking three consecutive slices together. Using a sliding window, we generated *n*-2 such partial-3D-volumes from each MRI image that contains *n* slices. This also allows for a larger training data pool since each image slab is unique. On the other hand, three natural orientations—axial, sagittal, and coronal—are available for use. To avoid the bias from the three orientations and to provide additional robustness to artifacts that may exist in the data, we pooled all partial-3D-volumes from the three orientations together during the training process. So, an *n*×*n*×*n* MRI image can yield 3 × (*n* – 2) training samples. For each of the orientations, we generated a 3D volume by stacking all the partial-3D-volumes together. In this case, if one slice belongs to multiple partial-3D-volumes the output is generated by averaging the specific slice from all partial-3D-volumes. The final MRI volume is the average of the three 3D volumes generated using all three orientations. Furthermore, during the harmonization, we applied a slice-matched harmonization strategy which relies on the brain registration. That is, after the registration, the reference slice selected for harmonization lies in the same coordinates as the source slice to be harmonized.

### 2.2. Model Inputs

#### 2.2.1 Datasets

We obtained T1-weighted brain MR images from five publicly available datasets: UK Biobank (UKBB), Parkinson’s Progression Markers Initiative (PPMI), Alzheimer Disease Neuroimaging Initiative (adni.loni.usc.edu; The ADNI was launched in 2003 as a public-private partnership, led by Principal Investigator Michael W. Weiner, MD. The primary goal of ADNI has been to test whether serial magnetic resonance imaging (MRI), positron emission tomography (PET), other biological markers, and clinical and neuropsychological assessment can be combined to measure the progression of mild cognitive impairment (MCI) and early Alzheimer’s disease (AD)), Adolescent Brain Cognitive Development (ABCD) and International Consortium for Brain Mapping (ICBM). See acknowledgements for more information on datasets.

Scans used in this study were collected from subsets of disease-free participants (UKBB: n=200, age range 45-55 years old; PPMI: n=76 age range 67-70 years old; ADNI: n=42, age range 67-70 years old; ABCD: n=200, age range 9-11 years old; and ICBM: n=200, age range 19-54 years old), among which 90% were used as training/validation sets and 10% testing sets. To ensure we can validate the disentangling of style from biological variables, we kept some cohorts overlapping in age, a demographic variable with very large effects. This way, the styles extracted from the cohorts overlapping in age can be compared directly without concern for whether they were determined by major biological factors, specifically age. As in most organic cases, the number of scans per dataset was not kept equal. All image acquisition information for these public resources can be found elsewhere, but briefly they vary in terms of scanner manufacturer, field strength, voxel size, and more, often within the same study. A list of manufacturers and field strength of the five datasets used in our study can be found in **Table 1**.

**Table 1.** Scanner manufacturer, field strength and age range specifications for the 5 dataset subsets used for training the model. The selection of images used here might not be representative of the entire dataset.

| Datasets | Scanner manufacturers |  |  |  | Field strength |  |  | Number of images | Age range (years) |
| --- | --- | --- | --- | --- | --- | --- | --- | --- | --- |
|  | GE | PHILLIPS | SIEMENS | ELSCENT | 1.5T | 2T | 3T |  |  |
| ADNI3 |  |  | ✓ |  |  |  | ✓ | 42 | 67-70 |
| ICBM | ✓ | ✓ |  | ✓ | ✓ | ✓ | ✓ | 200 | 19-54 |
| UKBB |  |  | ✓ |  |  |  | ✓ | 200 | 45-55 |
| PPMI | ✓ | ✓ | ✓ |  | ✓ |  | ✓ | 76 | 67-70 |
| ABCD | ✓ | ✓ | ✓ |  |  |  | ✓ | 200 | 9-13 |

#### 2.2.2 Image Processing

Many image processing steps are well established, can be implemented quickly, and can help reduce unnecessary variability in site differences that might be attributable to style. Rather than start from raw MRI images, all the images were skull-stripped using HD-BET [31], nonuniformity corrected using N3 approach, and linearly registered to the 1 mm^3^ MNI152 template using FSL *flirt* (9 degrees of freedom). The images were then resized to 0.8 mm^3^ isotropic 256×256×256 voxels to help prepare for 3D processing in all orientations.

### 2.3. Experimental Datasets for Model Evaluation

We evaluated our harmonization model on images that were not included in the training set, but were part of the datasets used in training (see details in Table 2). To make sure the images did not have major biological differences, we selected MR images from healthy subjects who were scanned between 55 age and 65 years from three datasets: UKBB (n = 300; age=60.06 ±2.96 year old); PPMI (n = 185; age=59.68 ±3.17 years old); and ADNI (n = 290; 60.09 ± 2.55 years old). To test whether the harmonization would over-correct the pathological alterations, we further harmonized scans from ADNI participants diagnosed with dementia within the same age range (n = 350; age=59.96 ±2.74 years old) referenced by a random healthy scan from the UKBB. Within the ADNI dataset, we tested whether dementia versus control differences remained consistent after harmonization.

**Table 2.** Datasets used for each of the validation experiments. Testing images denote the images are from the 10% testing dataset listed in Table 1. Extra images denote the images are not included in **Table 1**.

|  |  |  | Validation experiments |  |  |  |  |  |  |
| --- | --- | --- | --- | --- | --- | --- | --- | --- | --- |
|  |  |  | Histogram comparison | Intra-inter subject comparison | t-SNE | Cortical features comparison | Brain age prediction | Traveling subjects (1.5T vs. 3T) | Traveling subjects (12 scans) |
| Datasets | Testing images | ADNI | N=4<br>Age range=67-70 |  |  |  |  |  |  |
|  |  | PPMI | N=8<br>Age range=67-70 |  |  |  |  |  |  |
|  |  | UKBB |  | N=10<br>Age range=45-55 |  |  |  |  |  |
|  | Extra images | ADNI |  |  | N=290<br>Age range=55-65 | N=290<br>Age range=55-65 | N=220<br>Age range=55-75 | N=44<br>Age range=55-72 |  |
|  |  | PPMI |  |  | N=185<br>Age range=55-65 | N=185<br>Age range=55-65 | N=190<br>Age range=47-76 |  |  |
|  |  | UKBB |  |  | N=300<br>Age range=55-65 | N=300<br>Age range=55-65 | N=200<br>Age range=47-78 |  |  |
|  |  | Tong |  |  |  |  |  |  | N=3<br>Age range=23-26 |

To further validate our harmonization model, we applied the trained harmonization model to two traveling subjects datasets. The first dataset is a select portion of the ADNI-1 dataset: 44 subjects scanned with both a 1.5T scanner and a 3T scanner within 30 days of each other. We note, no scans from the 1.5T ADNI-1 dataset were used in model training. This dataset was used to test whether our model can harmonize the images from the same subjects scanned at different field strength MRIs to extract the same metrics. The second is the data described in [4], where three subjects were scanned twelve times at ten different sites within 13 months. Among the twelve scans, nine were at unique sites, and the remaining three scans were at the same site for every subject. The acquisition protocols were strictly controlled for all scan sessions.

### 2.4. Evaluation Metrics

The following evaluations were conducted on the experimental dataset, to compare datasets before and after the harmonization:

#### 2.4.1 Histogram comparisons

To quantitatively compare the intensity histogram of the harmonized images and reference images, we select all test participants from ADNI, and harmonize them with all test participants (age and sex matched) from PPMI. Participants were matched for age and sex to ensure approximately similar volumes of each tissue type. We compared the histograms of all paired brain-extracted MR images (one source ADNI scan and one target PPMI scan), and the histogram of the image after the ADNI scan was translated to the PPMI domain, using Jensen-Shannon (JS) divergence.

#### 2.4.2 Intra and inter subject image similarity

Ideally, MR image harmonization methods should not only remove the cross-sites variances, but also rigorously maintain the anatomical information within subjects. To test whether the anatomical information in MR images were preserved after harmonization, we compared the intra-subject similarity and inter-subject differences across the harmonizations. We selected 10 random subjects from the UKBB testing set as source images, and 100 randomly selected images from other datasets as reference images. That is, for each subject, we generated 100 harmonized images.

To determine how well the intra-subject similarity was preserved, we compared the similarity between images from the same subject across harmonizations, and the similarity of the images between pairs of subjects harmonized using identical reference images. The similarity was measured using intensity correlation (r), the structural similarity index measure (SSIM), and/or the peak signal-to-noise ratio (PSNR) as metrics.

To determine whether the inter-subject differences were also preserved, we first quantified the inter-subject differences by using the intensity differences to compute a Euclidean distance [5] between any two scans, forming a distance matrix, denoted as 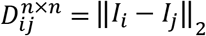, where n = 10 is the number of scans, and *I* the whole-image voxel intensity vectors for scans *i* and *j*. The goal was to estimate how the distances were preserved relative to each other before and after harmonization. We computed the correlation r between the two distance-matrices (only upper triangle) before and after harmonization.

#### 2.4.3 T-distributed stochastic neighbor embedding (t-SNE) of style-codes

To illustrate whether the 1×64 style code was successfully injected into the harmonized images, a t-distributed stochastic neighbor embedding (t-SNE) plot [32] was used to visualize the style representations of images randomly selected from the ADNI, UKBB and PPMI datasets, respectively. Briefly, t-SNE is a non-linear dimensionality reduction method for visualizing high dimensional data, where more similar data points are closer together, and dissimilar points further apart. The style code was extracted from the style encoder trained in the model before and after the images were harmonized.

### 2.5 Task-Specific Evaluation Analyses

#### 2.5.1 Comparisons of FreeSurfer derived cortical and subcortical measures

Cortical surface reconstruction and subcortical volume segmentation were performed using the freely available FreeSurfer 7.1.0 image analysis software (http://surfer.nmr.mgh.harvard.edu/)[33]. The bias field correction and skull-stripping were skipped in the recon-all script because these steps were performed as part of the image preprocessing steps before harmonization. Volumes in regions of interest (ROIs) were obtained directly from FreeSurfer’s aseg.stats and aparc.stats files. Thicknesses of cortical ROIs were obtained from FreeSurfer’s output *.aparc.stats files. Internally, FreeSurfer’s cortical thickness algorithm calculates the mean distance between vertices of a corrected, triangulated estimated GM/WM surface and GM/CSF (pial) surface [34].

Our FreeSurfer features of interest included: lateral ventricle volumes, hippocampal volumes, cerebral gray matter and cerebral white matter volume, and cortical gray matter thickness. These features were used to compare case/control effect sizes before and after harmonization, and used to determine cross-site similarities in extracted feature values for subjects scanned across multiple scanners.

#### 2.5.2 Brain Age

Brain age is a relatively new concept aimed at providing the age that would be predicted for an individual given their brain MRI scan. In older individuals, a brain age prediction higher than the individual’s true age may suggest accelerated aging [35]. We used a deep-learning based brain age prediction model as in [36] to predict the brain age before and after the harmonization. The brain age prediction model applied in this study takes a 3D scan as input and encodes each slice using a 2D-CNN encoder. Next, it combines the slice encodings using an aggregation module, resulting in a single embedding for the scan. Finally, this embedding is passed through the feed-forward layers to predict the brain age. The model is trained end-to-end using MSE loss. An overview of the network architecture of the brain age prediction model can be found in [36].

Here, the brain age prediction model was trained using the brain MR images of 1400 healthy UKBB participants between the ages of 45 and 76 years old. After the model was trained, it was applied to the T1-weighted images of a separate set of healthy subjects from the UKBB (n = 200; age range 47.3-77.4 years old), ADNI (n = 220; age range 55.6-74.8 years old), and PPMI (n = 190, age range: 47.2-75.5 years old) datasets. We then harmonized the ADNI and PPMI images using a reference image randomly selected from the UKBB dataset and applied the brain age prediction model for a pre- and post-harmonization comparison of predicted brain age.

### 2.6 Traveling Subjects Evaluation Analyses

#### 2.6.1 Traveling subjects who were scanned on 1.5T and 3T scanners

To provide a ground truth, we applied our harmonization model on two traveling subjects cohorts. One is from the ADNI dataset, who were scanned on 1.5T and 3T scanners. FreeSurfer metrics were extracted from all the images and compared between 1.5T images and 3T images before and after the harmonization. The percent volume differences (delta volume) were calculated by dividing the volume differences values by the average of the two volumes (average of 1.5 T and 3T), and then compared. To prove the effectiveness of our method, we further harmonized the structural volume using a classic harmonization method for image derived features, namely ComBat [37], and then compared the volume differences between ComBat and our method.

#### 2.6.2 Traveling subjects who were scanned from ten sites

For the evaluation on the second traveling subjects cohort from Tong et al. [4], we highlight how the reference can be to an image from a dataset not in the initial model training. One randomly selected image from the PPMI dataset was chosen as the reference image to harmonize all other MR images for all subjects. The PSNR and SSIM were measured between all pairs of images for the same subjects for comparisons between images before and after harmonization.

### 2.7 Comparisons with Other Harmonization Networks

We trained the other two models with two state-of-the-art unsupervised harmonization methods, namely cycleGAN and starGAN, and compared them with our method. The images of 10 ADNI subjects were harmonized to a single UKBB subject. Input and output images were subtracted to compare preservation of anatomical structure.

## 3. Results

### 3.1 Hyperparameter Tuning

Figure 2 shows an example of an image harmonized using the same model but with different *λ_cyc_* values. In this example, the source image is from ADNI and the reference image is from ABCD. There is a 58-year age difference between these subjects, and very evident differences in anatomical structure, including larger lateral ventricles in the older adults. If *λ_cyc_* = 0, meaning none of the style-irrelevant characteristics are needed, the model learns everything from the reference image, generating an image completely identical to the reference. If *λ_cyc_* = 1, then the model learns the style from the reference image but also some biological patterns, such as smaller lateral ventricles and thicker gray matter cortices. If *λ_cyc_* = 10, then the model learns only the style information from the reference and rigorously maintains the style-irrelevant characteristics (i.e., ventricle and other regional volumes) from the source images.

**Figure 2.**
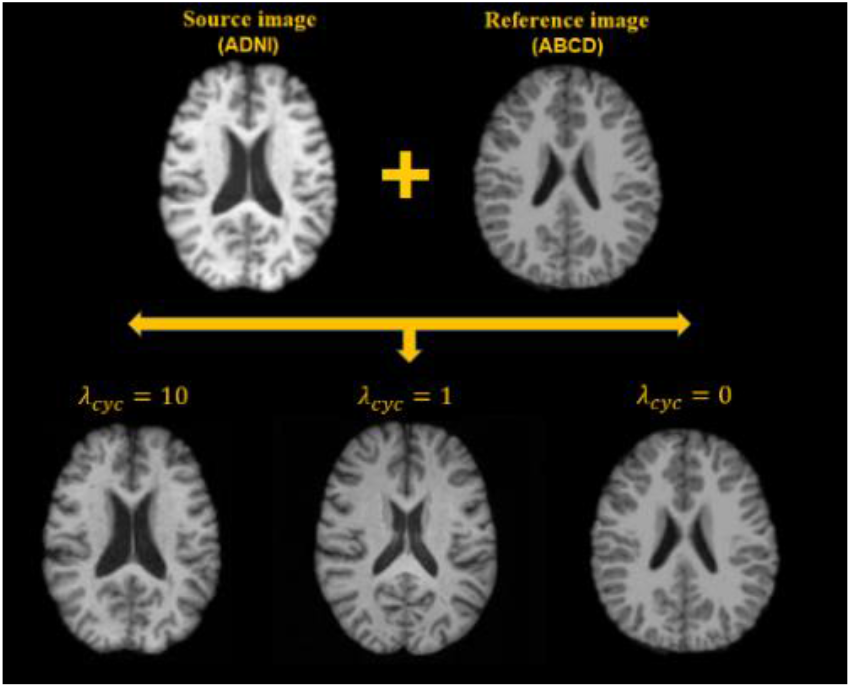
Small cycle consistency loss coefficients bias the generated images towards the reference images, while larger cycle consistency coefficients rigorously maintain the structure of the source images, only altering the style.

### 3.2 Image-wide Evaluations

#### 3.2.1 Comparisons across cohorts before and after harmonization

Figure 3 illustrates the harmonized images among the five datasets according to nine randomly selected reference images from across the datasets. Qualitatively, harmonized images are noticeably more similar in contrast and intensity to those of the reference images, and their anatomical structures were well-maintained. In a quantitative comparison of the age- and sex-matched participants of the ADNI and PPMI datasets, the JS divergence between the histograms of the ADNI and the translated ADNI → PPMI image (0.047 ± 0.005) was significantly higher than that of the PPMI and the ADNI → PPMI translated image (0.023±0.006; p<0.0001), suggesting the histograms of the ADNI image harmonized to PPMI has an average intensity profile more similar to PPMI than ADNI, from which it came.

**Figure 3.**
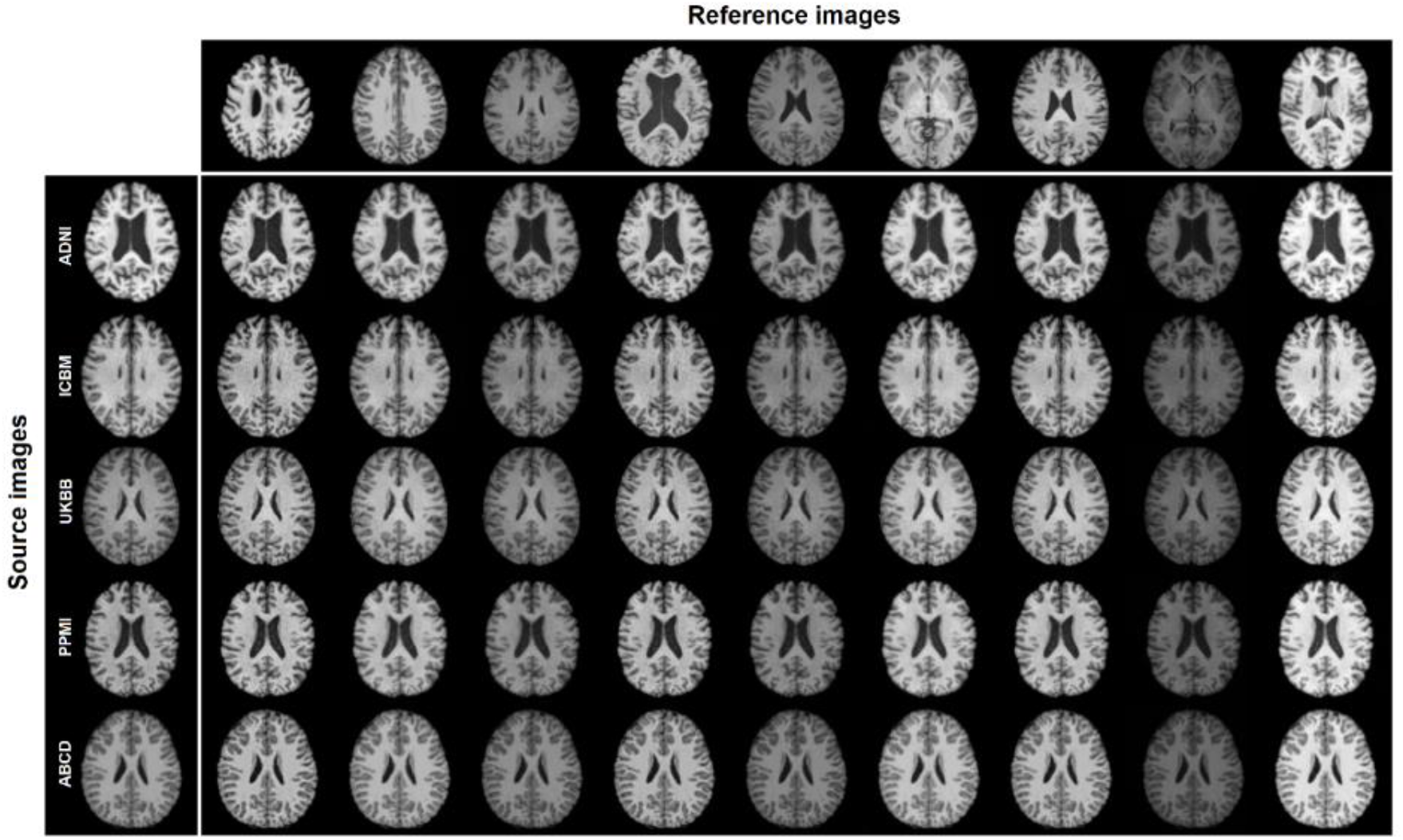
The style-encoding GAN can harmonize images based on a single reference image.

Our t-SNE results of the style representations reveal that before harmonization, the style features produced are separable by datasets, especially the PPMI dataset. After the harmonization the style features become jointly embedded and the style feature embedding is not informative of datasets (**Figure 4**), suggesting the effectiveness of the proposed method.

**Figure 4.**
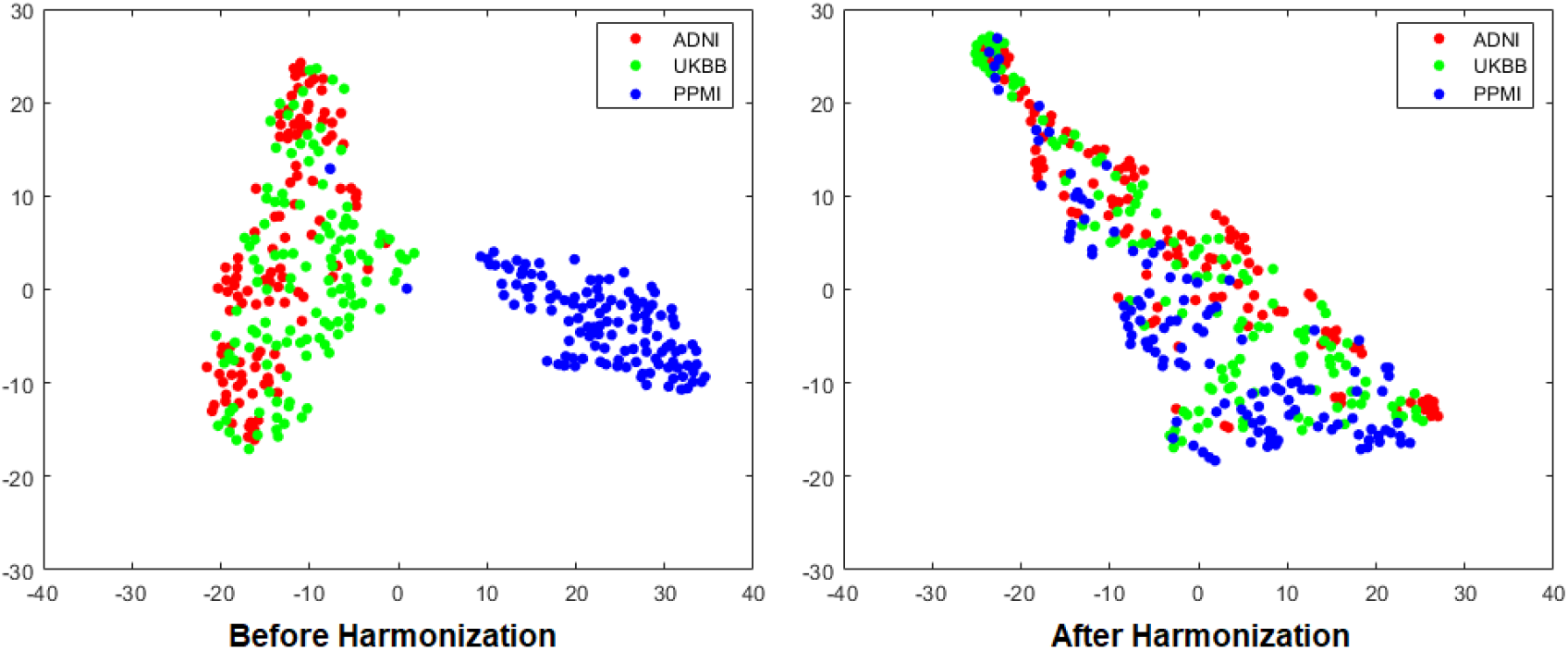
T-SNE representation of the style code extracted in images from three datasets (ADNI, UKBB, and PPMI) before and after the harmonization.

#### 3.2.2 Intra-subject similarity

When testing the intra-subject similarities between UKBB source subjects and reference subjects from other datasets, the average correlation between the intensity of the images from the same subjects across harmonizations was 0.991 and the average SSIM was 0.801. The average correlation between the intensity of the images from two different subjects in identical harmonization were r=0.889 and SSIM=0.517. For both sets of values, the intra-subjects similarities after different harmonizations were significantly higher than the inter-subject similarities after identical harmonizations, indicating the intra-subject anatomical information was preserved after the harmonization compared to inter-subject variances.

#### 3.2.3 Inter-subject differences

When computing the correlation *r* between the two image-to-image wide Euclidean distance-matrices as described, before and after harmonization, our model achieved an average correlation of *r*=0.979 (range: [0.954, 0.994]) between the distance-matrices before harmonization and the 100 distance-matrices after harmonization, indicating the inter-subject difference was reliably preserved after harmonization.

### 3.3 Task-Specific Evaluation of Downstream Analyses on 3D Reconstructions

#### 3.3.1 FreeSurfer Cortical Thickness and Regional Volumes

We illustrated the downstream applications of our harmonization method using cortical features extracted from automated processing software, in this case, FreeSurfer. Cortical thickness measures were generated from healthy subjects aged between 55 and 65 years in three datasets (UKBB, ADNI and PPMI). Volumes from several brain structures and cortical thickness distributions from left and right hemispheres are plotted in **Figure 5** and the values are shown in **Table 3**, and the p-values from the two-tailed t-tests between pairs of datasets are shown in **Table 4**. Several features were significantly different between datasets before harmonization, including, for example, the total cortical gray matter volumes between ADNI and UK Biobank; after harmonization of these images, no statistical differences were detected between the GM volumes of these datasets. Overall, after harmonization, the brain structural volumes and cortical thickness estimates extracted and evaluated here from the three datasets showed distributions with means that were not statistically different after harmonization Table 4).

**Figure 5.**
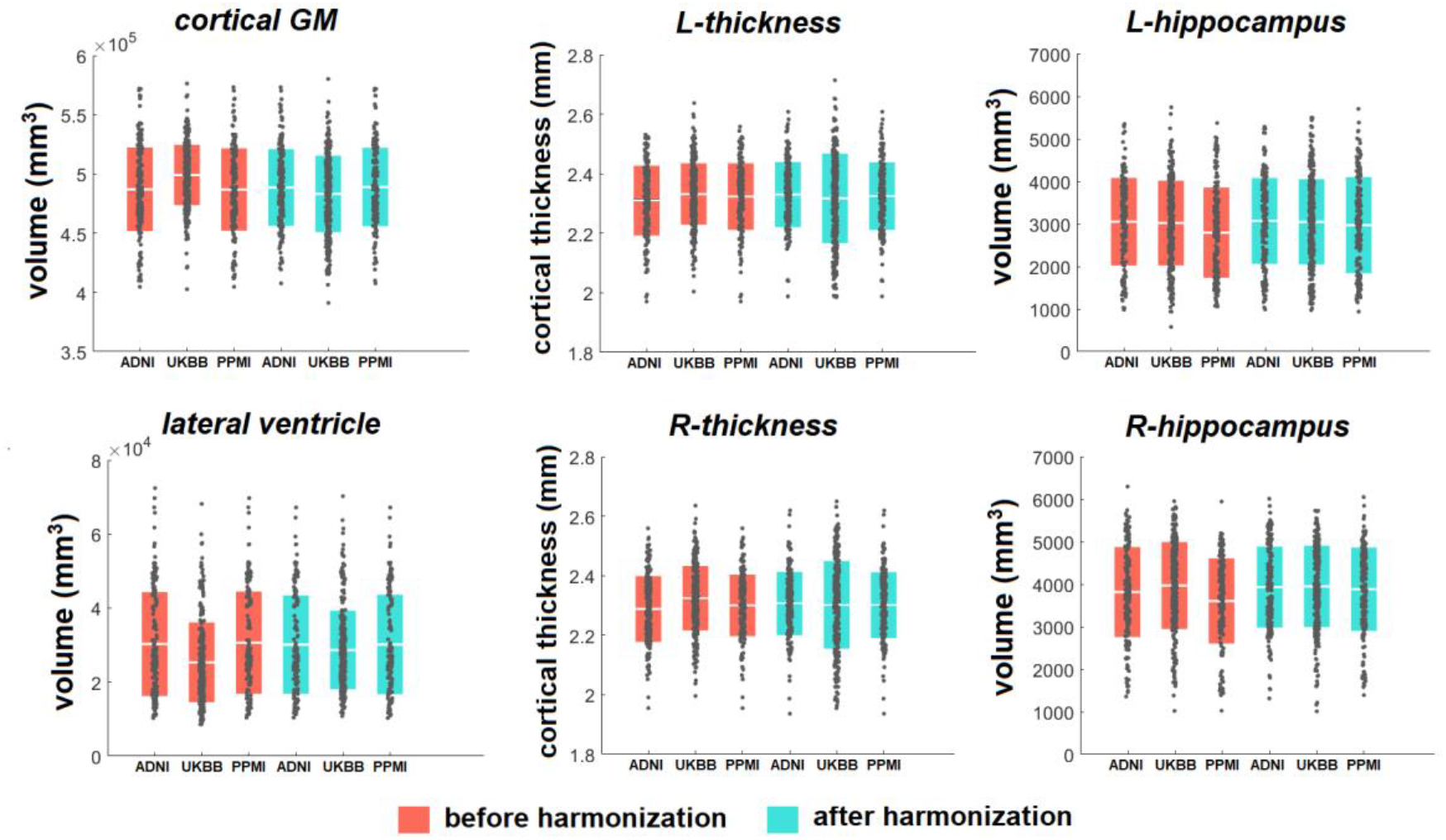
Brain structural volumes and cortical thickness comparisons among the three datasets before and after the harmonization.

**Table 3.** Brain structural volumes and cortical thickness among the three datasets before and after harmonization.

| | Cortical GM<br>($10 \times 10^5 \text{ mm}^3$) | | Lateral<br>ventricle<br>($10 \times 10^4 \text{ mm}^3$) | | L GM<br>thickness<br>(mm) | | R GM<br>thickness<br>(mm) | | L<br>hippocampus<br>($\text{mm}^3$) | | R<br>hippocampus<br>($\text{mm}^3$) | |
| --- | --- | --- | --- | --- | --- | --- | --- | --- | --- | --- | --- | --- |
|  | before | after | before | after | before | after | before | after | before | after | before | after |
| ADNI | 4.84<br>$\pm 0.49$ | 4.88<br>$\pm 0.32$ | 3.01<br>$\pm 1.41$ | 3.02<br>$\pm 1.32$ | 2.31<br>$\pm 0.13$ | 2.34<br>$\pm 0.11$ | 2.29<br>$\pm 0.13$ | 2.33<br>$\pm 0.11$ | 3055<br>$\pm 1030$ | 3070<br>$\pm 1008$ | 3816<br>$\pm 1057$ | 3932<br>$\pm 953$ |
| UKBB | 4.99<br>$\pm 0.28$ | 4.83<br>$\pm 0.32$ | 2.60<br>$\pm 1.25$ | 2.89<br>$\pm 1.23$ | 2.34<br>$\pm 0.12$ | 2.33<br>$\pm 0.15$ | 2.34<br>$\pm 0.12$ | 2.32<br>$\pm 0.15$ | 3021<br>$\pm 996$ | 3050<br>$\pm 1003$ | 3969<br>$\pm 1021$ | 3941<br>$\pm 956$ |
| PPMI | 4.85<br>$\pm 0.34$ | 4.89<br>$\pm 0.32$ | 3.04<br>$\pm 1.39$ | 3.01<br>$\pm 1.38$ | 2.33<br>$\pm 0.13$ | 2.33<br>$\pm 0.12$ | 2.31<br>$\pm 0.12$ | 2.32<br>$\pm 0.12$ | 2794<br>$\pm 1062$ | 2974<br>$\pm 1132$ | 3605<br>$\pm 1001$ | 3879<br>$\pm 980$ |

**Table 4.**
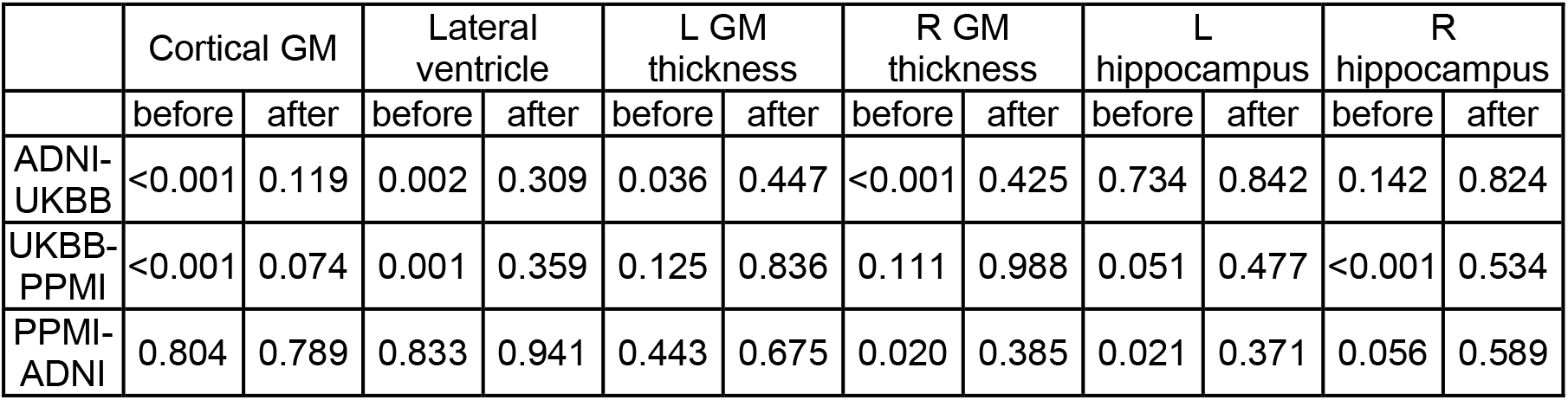
The p-values of two-tailed t-test of brain structural volumes and cortical thickness between pairs of the datasets before and after the harmonization.

#### 3.3.2 Maintaining *Case/Control Effect Size Differences*

A key obstacle for a good harmonization method is to avoid the over-correction for meaningful biological features, especially those which may help distinguish between pathological groups and healthy control groups. To illustrate that the clinical features can be well-preserved after harmonization using our method, we compared the hippocampal volumes between AD patients and healthy subjects in the ADNI dataset, before and after the harmonization to a UKBB reference. Hippocampal volume was extracted for left and right hemispheres respectively. Before harmonization, the left and right hippocampal volumes in AD patients (left: 2127.9 ±985.8 *mm*^3^, right: 2676.2.9 ± 1026.6 *mm*^3^) were significantly smaller than that of healthy subjects (left: 3130.7.6 ± 1009.1 *mm*^3^; right: 3835.6 ± 1023.8 *mm*^3^; left: p < 0.0001, Cohen’s d = −0.98; right:p < 0.0001, Cohen’s d = −1.14). As depicted in Figure 9, these differences remained robust after harmonization as well; the hippocampal volumes in AD patients (left: 2298.6 ± 1010.1 *mm*^3^; right: 2858.6 ± 923.5 *mm*^3^) compared to controls (left: 3129.9 ± 1008.3 *mm*^3^, right: 3930.2 ± 953.0 *mm*^3^) remained significant with effect size differences nearly identical to those before harmonization (left: p < 0.0001, Cohen’s d = −0.83, right: p < 0.0001, Cohen’s d = −1.13) (**Figure 6**).

**Figure 6.**
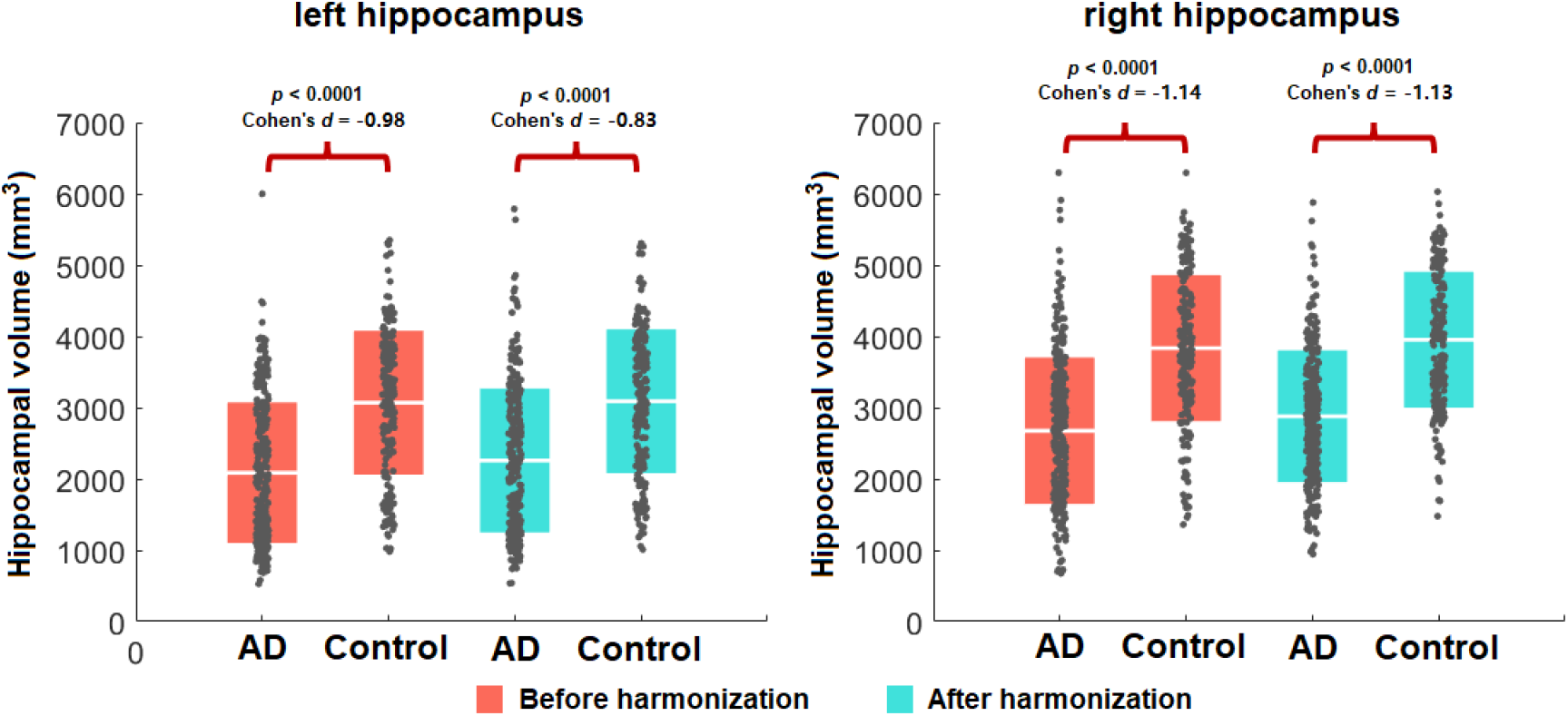
Hippocampal volume comparisons among participants with an AD diagnosis to cognitively healthy controls within the ADNI dataset, before and after the harmonization. Harmonization does not affect the within-cohort statistical case/control differences.

#### 3.3.3 Brain Age Prediction

In the UKBB healthy brain age test set, a mean absolute error (MAE) of 3.47 years was achieved between the true chronological age and the predicted brain age, and the Pearson correlation coefficient was 0.82. We observed poor generalization ability to other datasets before harmonization (for ADNI: MAE = 4.93 years, Pearson correlation coefficient = 0.45; for PPMI: MAE = 4.77 years, Pearson correlation = 0.79). After harmonization of images from ADNI and PPMI to a reference image from UKBB, we found an improvement in the generalization performance of our predictor (for ADNI: MAE = 3.79 years, Pearson correlation coefficient = 0.58; and for PPMI: MAE = 3.88 years, Pearson correlation = 0.84; **Figure 7**).

**Figure 7.**
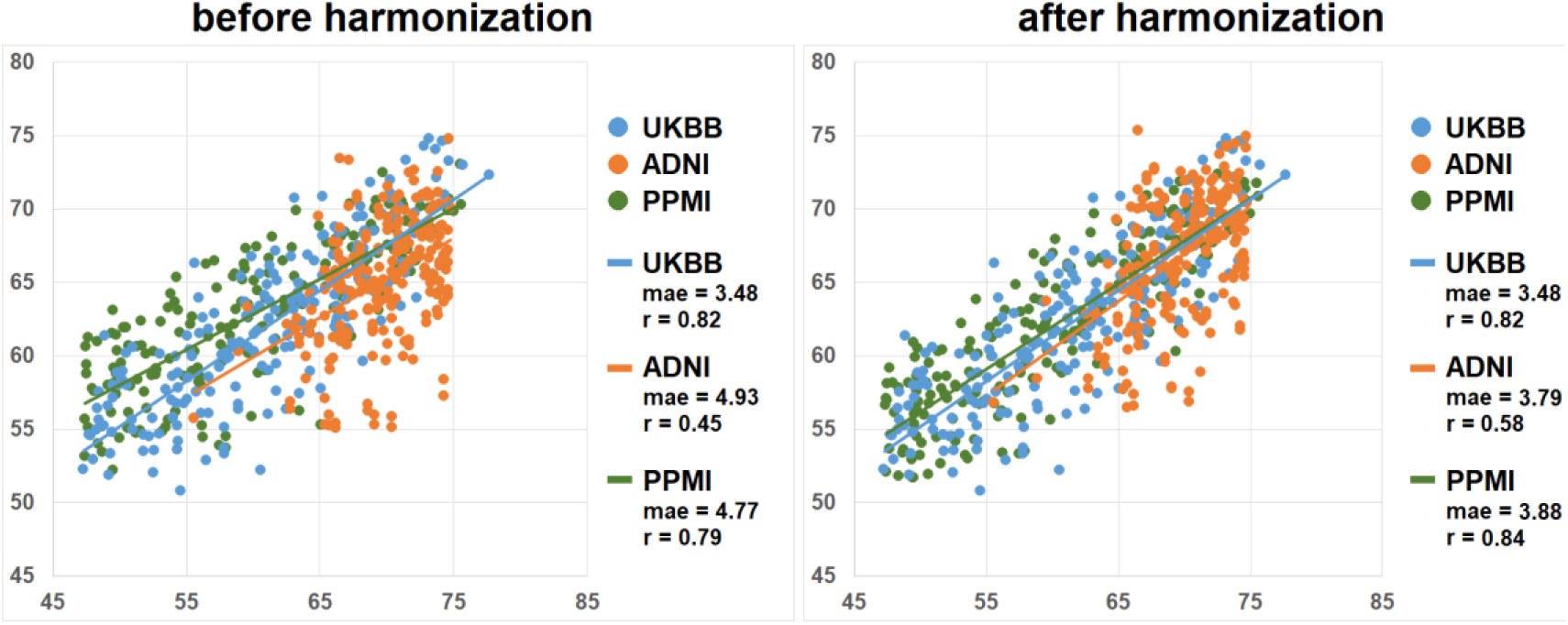
Brain age prediction comparisons among the three datasets before and after the harmonization. The brain age prediction model was trained using 1400 healthy scans from the UKBB dataset. Before harmonization, the mean absolute errors (MAEs) were much lower when the test set was from the same data collection site as the training data than other datasets, but after harmonization, the errors from the other cohorts were minimized.

### 3.4 Traveling Subjects Evaluations

#### 3.4.1 Harmonization of Traveling Subjects Scanned on 1.5T and 3T Scanners

After harmonization, volume differences between 1.5T image and 3T images were smaller than before harmonization for all the brain structures we evaluated. Two-sided t-tests indicate that after harmonization, volume difference between 1.5T and 3T images is significantly smaller than before harmonization for hippocampal volume (p = 0.002). comparisons of the volume differences between ComBat and our method showed that our method outperformed the ComBat for significantly smaller volume difference between 1.5T and 3T for hippocampal volume (p = 0.004). Our method also exhibited comparable effects as ComBat for other brain structures (p’s > 0.21) in this application (**Figure 8**).

**Figure 8.**
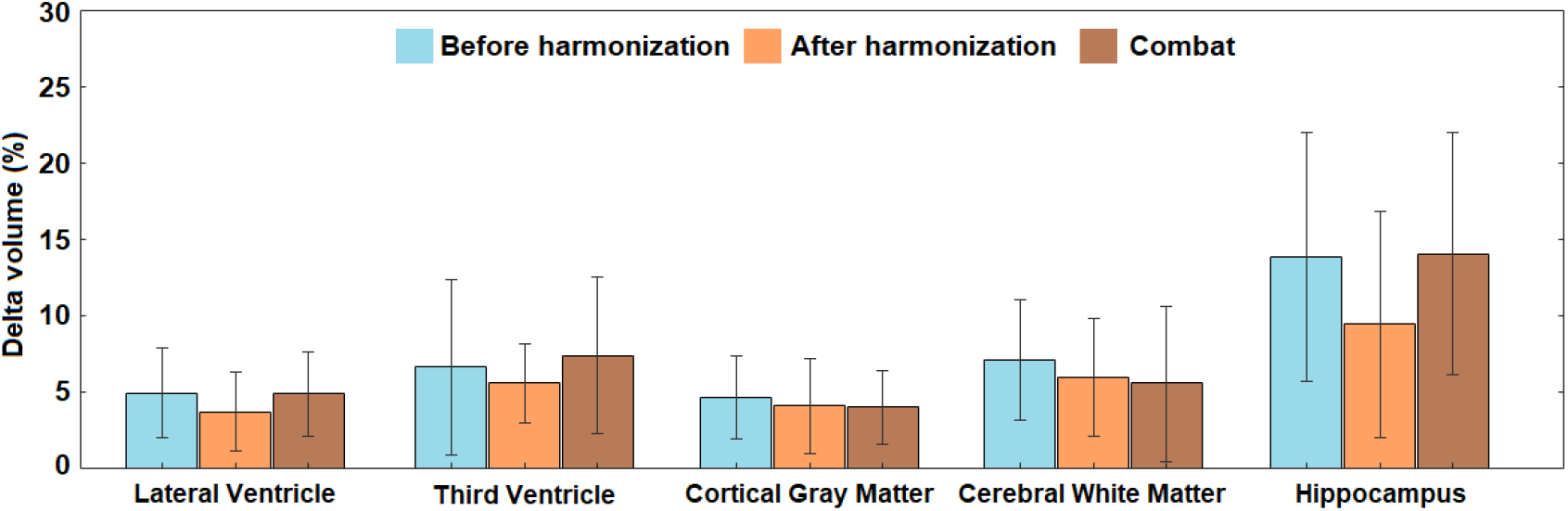
Cortical structural volume differences for traveling subjects scanned by 1.5T scanner and 3T scanner within 30 days. The comparison was made among the volume differences before the harmonization, after the harmonization using our method, and after harmonization using combat.

#### 3.4.2 Harmonization of Traveling Subjects from ten Sites

In the traveling subjects cohort from Tong et al. [4], we highlight how the reference can also be to an image from a dataset not in the initial model training. One randomly selected image from the PPMI dataset was chosen as the reference image to harmonize all other MR images for all subjects. The PSNR and SSIM were measured between all pairs of images for the same subjects. That is, for each subject, we have 45 pairs of cross-site pairs (from 9 scans) and 6 pairs of same-site pairs (from 3 scans). The average and standard deviation values for each subject are shown in **Figure 9**, which shows that the harmonized images, either for cross-sites scans and same-site scans, are more similar in appearance. Quantitative results for the traveling subjects show a dramatic improvement in similarity using both SSIM (0.954 for original images vs. 0.969 for harmonized images) and PSNR (M = 26.1 for original images and M = 28.2 for harmonized images), two-sided t-tests, p’s < 0.01.

**Figure 9.**
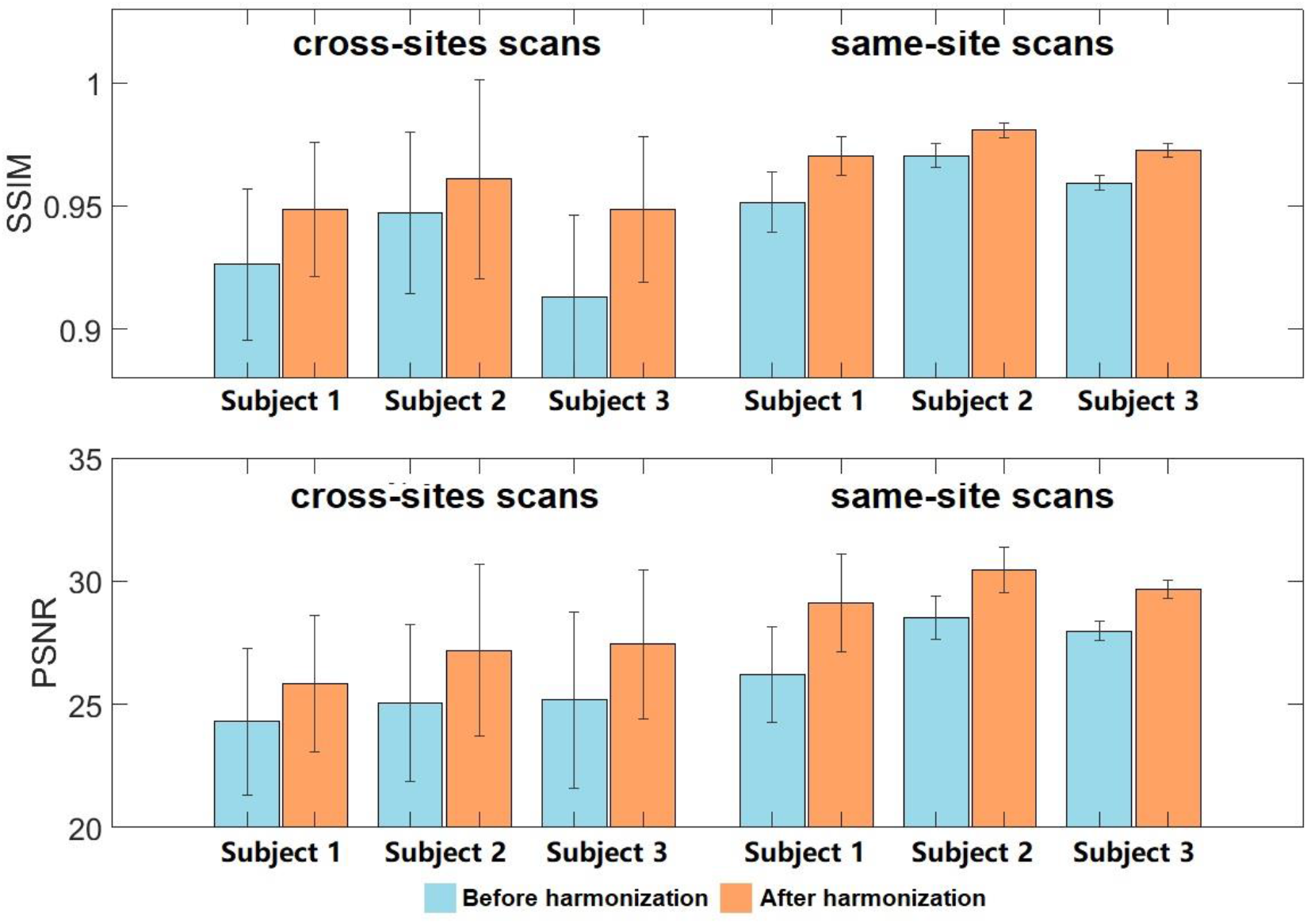
Image similarity comparisons using the structural similarity index (SSIM) and peak signal-to-noise ratio (PSNR) for three traveling subjects, each scanned 12 times at 10 different sites, all within 13 months.

We further evaluated our model by harmonizing images from the unseen dataset (one randomly selected image from the traveling subjects in [4] to the images in our training datasets. **Figure 10** shows, qualitatively, that our model successfully captures the styles of the unseen/traveling subject and renders these styles correctly to the source images.

**Figure 10.**
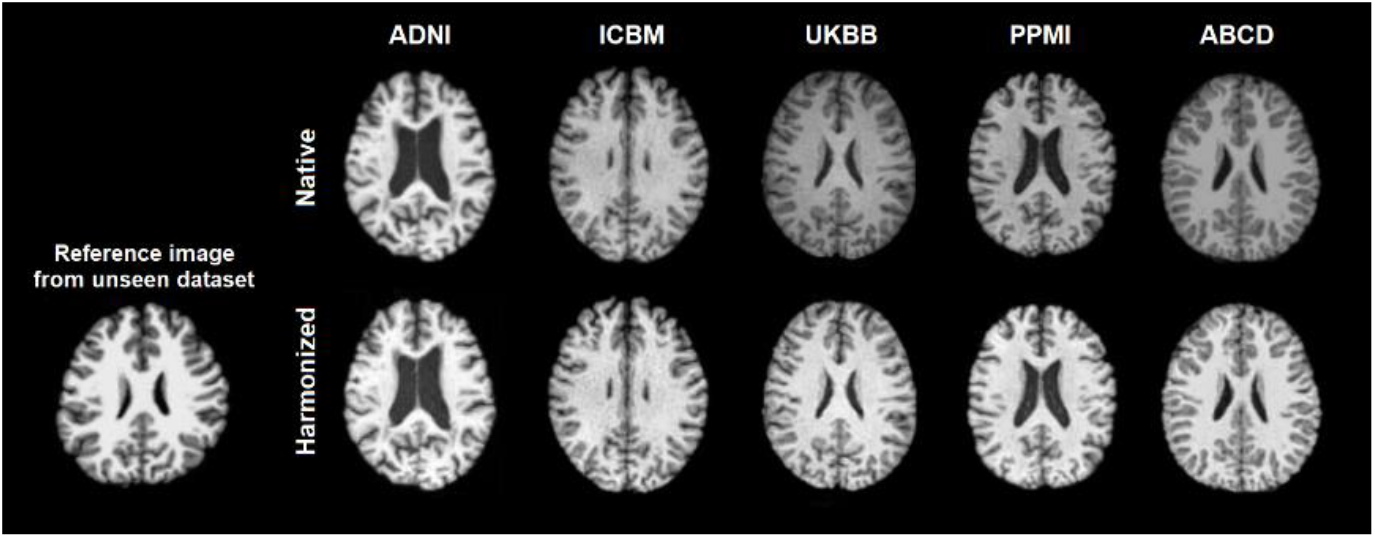
The trained style-encoding GAN successfully captures styles of reference images from novel acquisition protocols and renders these styles correctly to the source images.

### 3.5 Comparisons with other harmonization network structures

A visual inspection of harmonization from ADNI dataset to UKBB dataset can be found in **Figure 11**. The results revealed that both cycleGAN and starGAN have some deficit in brain ventricles, which is not found in our method.

**Figure 11.**
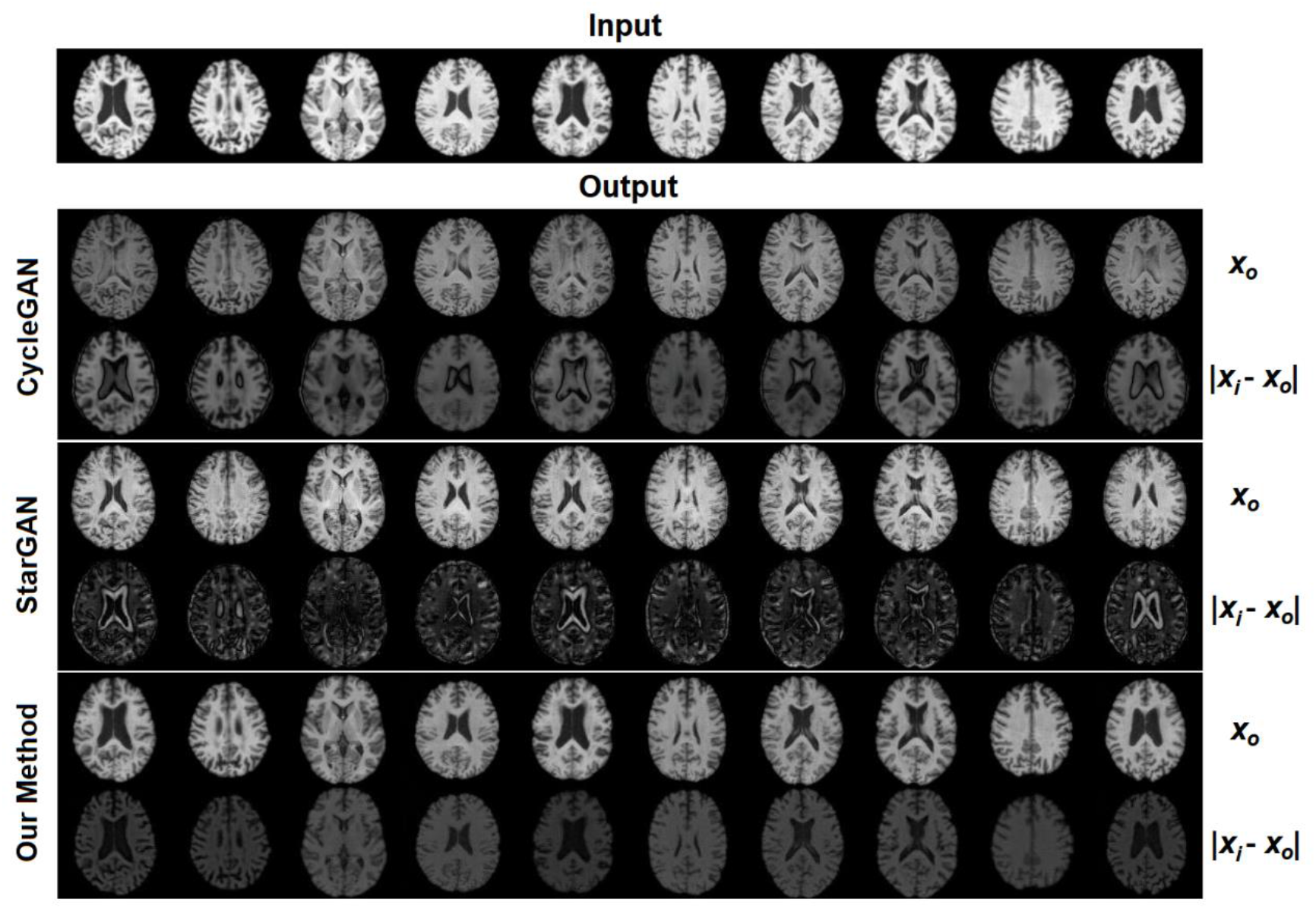
Visual comparison across three methods (CycleGAN, StarGAN, our Style Encoding GAN) of image harmonization using 10 images from subjects within the ADNI dataset, harmonized to a single subject within the UKBB dataset. For each method, the first row is the harmonized images translated from the input, and the second row is the absolute difference between the input and the harmonized images. Anatomical structure is not preserved in CycleGAN or StarGAN as specifically evident by the alterations within the ventricles and other contrast differences around tissue boundaries.

## 4. Discussion

We have developed a novel harmonization approach for T1-weighted MRIs collected across multiple vendors and acquisition protocols using a style-encoding GAN. In this work, a style-harmonized and anatomically preserved image is synthesized from a single input image from an arbitrary site, taking the style code encoded from an arbitrary reference image (usually from another site) directly as extra information, without knowing details of the acquisition protocols *a priori*. Our model does not rely on traveling subjects or any paired modalities of images or any other paired information from the same subjects. Furthermore, because we consider the cross-site harmonization as a style transfer problem rather than a domain transfer problem, the MR images from multiple sites do not need to be categorized into specific domains (i.e, acquisition protocols, specific scanners, studies, clinical conditions, age bins, etc). Thus, the demographic and pathological conditions do not need to be matched for the harmonization.

Unlike other harmonization approaches that work on harmonizing image derived features, and prepare harmonized features for specific tasks, our method works directly to harmonize the full brain MRI, from which already harmonized features can then be extracted. We have tested our method on several tasks including automatic image segmentation software, brain age predictions, and case/control effect size calculations. All of these applications highlighted the effectiveness of our model.

Most of the current image-to-image harmonization methods translate images between different domains, such as specific acquisition protocols or studies [15, 38]. However, “domain” itself is a complex concept. While it may seem straightforward to group images into different domains, such as the dataset or study where they come from, a study may oftentimes collect images across many sites, scanners, and acquisition protocols, suggesting more complex, nested domains. Even if the images were grouped as domains according to identical collection sites, scanners, or acquisition protocols, they may still exhibit within-domain variabilities, due to scanner drift over time as seen in the traveling subjects’ dataset from [4]. In short, the scenarios where domain-based approaches can optimally fit include a) if the image translation is conducted among limited groups with clear definitions, which is not realistic in many datasets, or b) when the domain is actually limited to a single specific image, as in our case. In other words, here, we do not separate images into domains based on datasets but consider every single image as a unique “domain” with its own style. While some methods make assumptions about the distribution of styles, for example that they match a universal prior, such as a Gaussian distribution, that spans all images and can be learned using a variational auto-encoder as in [21], our method does not make any such assumptions about the style distributions. We learn style codes adversarially using a GAN-like approach, which does not rely on any hypothesized prior distribution and allows us to learn style codes from every single image individually with greater flexibility and accuracy. An important caveat in biological data harmonization across data collection sites, including brain MRI harmonization, is that the biological information (i.e. brain anatomy) and non-biological information are convoluted, where demographic and clinical characteristics are often also dependent on the cohort, specifically the study inclusion and exclusion criteria. In these common instances, harmonization methods can easily “harmonize” both sets of information, leading to inadequate image harmonization and over-correction [11]. Disentangling images into content and style spaces can overcome this issue. Disentangled latent spaces have been used in several past image translation studies [20, 21]. These studies both extracted the brain structures as content explicitly, which requires an extra step to supervise the content learning using either different modality of images from the same subjects [20], or an extra content decoder [21]. To preserve the content, in this case being the anatomical information in the brain MRI, we propose not to generate such an explicit content code. We preserve the content information using a cycle-consistency loss by directly matching the source image and the image translated from the target image based on source style code. In this way, no extra paired information is needed, and we can still avoid the overcorrection of image acquisition confounded by biologically relevant information.

An important aspect of harmonization is, therefore, to keep the relevant biological and clinical patterns in the images without this overcorrection. We provide evidence that our model can preserve these patterns using the brain age prediction and hippocampal volume comparison in AD patients compared to age and sex matched controls. Brain age estimation has become an established biomarker of overall brain health in the neuroimaging community, exhibiting overlapping neuroanatomical patterns with a variety of other pathologic processes [39, 40]. Accurate brain age estimation depends on fine neuroanatomical patterns that can be obfuscated by cross-site imaging variations. Therefore, brain age is an excellent candidate experiment to assess harmonization performance. When training our model on one dataset, we demonstrated improved age prediction estimates in three separate sites, following their mapping to a reference image within the dataset used to train the brain age model. We further demonstrate how case control differences in hippocampal volumes between ADNI participants with dementia compared to cognitive healthy controls were not affected by the harmonization procedure. As the harmonization was to a reference image of a healthy individual, a harmonization procedure that would overcorrect, and confound imaging and biological sources of variability, would likely remove some of the anatomical variability due to the dementia and the case/control effect sizes would be smaller after harmonization than with the original data. That was not the case with our harmonization approach. In other words, we demonstrated that the degree of impairment captured by hippocampal volumes in the patient population was preserved after the harmonization, indicating the clinical patterns in different datasets were not over-corrected by our harmonization method.

Finally, we showed our model generalizes well even to unseen samples, effectively being able to harmonize to a reference image not included in any of the datasets used for model training. This is a particularly important advantage for studies with very small sample sizes, or those using less common acquisition protocols. For example, if investigators in charge of a relatively small, or unique, cohort wanted to compare their subjects with data from a large, open resource, they may use our method to avoid overcorrection and obtain similar styled-images.

Although the model was designed to separate brain structures (contents) and anatomy-irrelevant information (styles) completely, the model sometimes may not automatically and accurately recognize which are contents and which are styles. In other words, the styles recognized by the model may contain some of the content. This is controlled by the selection of hyperparameters, more experiments are needed to test how the hyperparameters may influence the harmonization. The outcome of the harmonization is dependent on the reference image, and here, we only tested reference images from individuals without gross brain structural abnormalities. It is possible content and style may be conflated in cases where the reference image may have severe artifacts, or anatomical abnormalities such as a large lesion. Our method works on T1-weighted images only, yet a similar framework may also be applied to harmonize images for other modalities, or the multi-modal image conversions.

In conclusion, here we develop a novel harmonization approach for T1-weighted MRIs using a style-encoding GAN that can be used to harmonize entire images for a variety of international, multi-cohort, neuroimaging collaborations.

## Data and Code Availability Statement

The data used in these experiments are available on application to the relevant studies. The code used is available when the paper is published and weights from training are available in USC-IGC/style_transfer_harmonization (github.com).

## Acknowledgements

This work was supported in part by: R01AG059874, RF1AG057892, U01AG068057, and P41EB015922. BrightFocus Research Grant award (A2019052S). This research has been conducted using the UK Biobank Resource under Application Number ‘11559’. Data used in preparation of this article were also obtained from the Alzheimer’s Disease Neuroimaging Initiative (ADNI) database (adni.loni.usc.edu). As such, the investigators within the ADNI contributed to the design and implementation of ADNI and/or provided data but did not participate in analysis or writing of this report. A complete listing of ADNI investigators can be found at: http://adni.loni.usc.edu/wp-content/uploads/how_to_apply/ADNI_Acknowledgement_List.pdf. Data used in the preparation of this article were also obtained from the Parkinson’s Progression Markers Initiative (PPMI) database (www.ppmi-info.org/data), the Alzheimer’s Disease Neuroimaging Initiative (ADNI) database (adni.loni.usc.edu), the Adolescent Brain Cognitive Development (ABCD) Study (https://abcdstudy.org), held in the NIMH Data Archive (NDA), and the traveling subjects cohort from Tong et al (https://www.nature.com/articles/s41597-020-0493-8#Sec7). For up-to-date information on the PPMI study, visit ppmi-info.org. PPMI – a public-private partnership – is funded by the Michael J. Fox Foundation for Parkinson’s Research and funding partners, including [list the full names of all of the PPMI funding partners found at www.ppmi-info.org/about-ppmi/who-we-are/study-sponsors].

Data collection and sharing for this project was funded by the Alzheimer’s Disease Neuroimaging Initiative (ADNI) (National Institutes of Health Grant U01 AG024904) and DOD ADNI (Department of Defense award number W81XWH-12-2-0012). ADNI is funded by the National Institute on Aging, the National Institute of Biomedical Imaging and Bioengineering, and through generous contributions from the following: AbbVie, Alzheimer’s Association; Alzheimer’s Drug Discovery Foundation; Araclon Biotech; BioClinica, Inc.; Biogen; Bristol-Myers Squibb Company; CereSpir, Inc.; Cogstate; Eisai Inc.; Elan Pharmaceuticals, Inc.; Eli Lilly and Company; EuroImmun; F. Hoffmann-La Roche Ltd and its affiliated company Genentech, Inc.; Fujirebio; GE Healthcare; IXICO Ltd.;Janssen Alzheimer Immunotherapy Research & Development, LLC.; Johnson & Johnson Pharmaceutical Research & Development LLC.; Lumosity; Lundbeck; Merck & Co., Inc.;Meso Scale Diagnostics, LLC.; NeuroRx Research; Neurotrack Technologies; Novartis Pharmaceuticals Corporation; Pfizer Inc.; Piramal Imaging; Servier; Takeda Pharmaceutical Company; and Transition Therapeutics. The Canadian Institutes of Health Research is providing funds to support ADNI clinical sites in Canada. Private sector contributions are facilitated by the Foundation for the National Institutes of Health (www.fnih.org). The grantee organization is the Northern California Institute for Research and Education, and the study is coordinated by the Alzheimer’s Therapeutic Research Institute at the University of Southern California. ADNI data are disseminated by the Laboratory for Neuro Imaging at the University of Southern California.

